# Modulating Mesenchymal Stromal Cell Microenvironment Alters Exosome RNA Content and Ligament Healing Capacity

**DOI:** 10.1101/2023.10.22.563485

**Authors:** Connie S. Chamberlain, Archana Prabahar, John A Kink, Erika Mueller, Yiyao Li, Stephanie Yopp, Christian M. Capitini, Peiman Hematti, William L. Murphy, Ray Vanderby, Peng Jiang

**Author notes:** To whom correspondence should be addressed: Peng Jiang. C.S.C and A.P contributed equally to this work.

## Abstract

Although mesenchymal stromal cell (MSC) based therapies hold promise in regenerative medicine, their applications in clinical settings remain challenging due to issues such as immunocompatibility and cell stability. MSC-derived exosomes, small vesicles carrying various bioactive molecules, are a promising cell-free therapy to promote tissue regeneration. However, it remains unknown mainly regarding the ability to customize the content of MSC-derived exosomes, how alterations in the MSC microenvironment influence exosome content, and the effects of such modifications on healing efficiency and mechanical properties in tissue regeneration. In this study, we used an *in vitro* system of human MSC-derived exosomes and an *in vivo* rat ligament injury model to address these questions. We found a context-dependent correlation between exosomal and parent cell RNA content. Under native conditions, the correlation was moderate but heightened with microenvironmental changes. *In vivo* rat ligament injury model showed that MSC-derived exosomes increased ligament max load and stiffness. We also found that changes in the MSCs’ microenvironment significantly influence the mechanical properties driven by exosome treatment. Additionally, a link was identified between altered exosomal microRNA levels and expression changes in microRNA targets in ligaments. These findings elucidate the nuanced interplay between MSCs, their exosomes, and tissue regeneration.

**Graphic Abstract:** 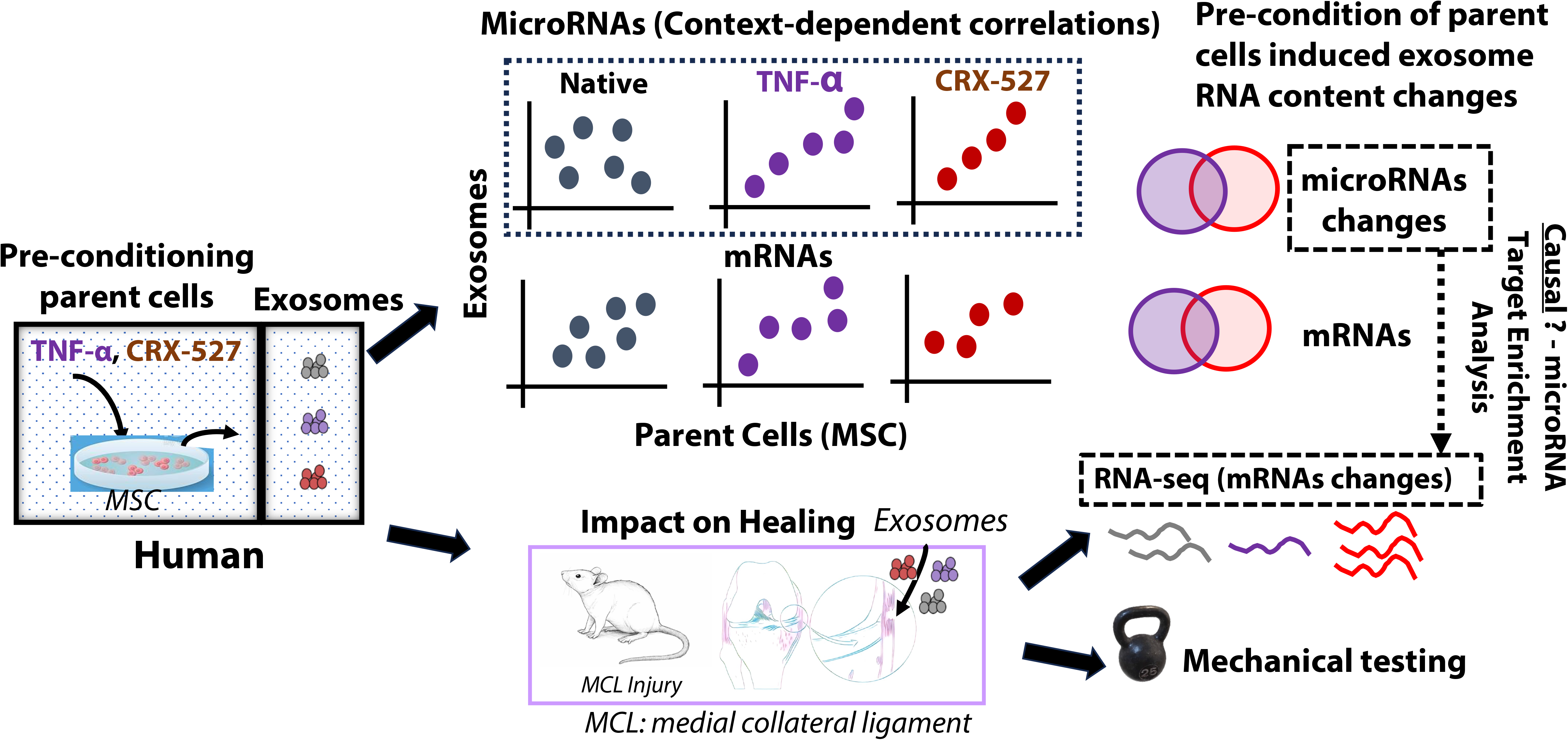

## Introduction

Mesenchymal stromal cells (MSCs), also known as mesenchymal stem cells, have emerged as a major focus for tissue regeneration. These multipotent cells hold immense potential for clinical applications due to their unique ability to promote tissue repair, immunomodulation, and regeneration^1^. The therapeutic benefits of MSCs stem from their capacity to release many growth factors and cytokines, which play a pivotal role in recruiting and activating other cells at the site of injury^2^. Moreover, MSCs facilitate the formation of new blood vessels, which is crucial for effectively healing damaged tissues^3^.

Despite the advancements in MSC-based therapies^4,5^, several challenges hinder their broad application in clinical settings. These challenges include but are not limited to immunocompatibility, stability, and migratory capacity^6^. Immunocompatibility remains a major concern as individual variations among donors and recipients may lead to immune responses and potential rejection of transplanted MSCs. Moreover, ensuring the stability of MSCs during storage, transportation, and transplantation is essential to maintain their regenerative potential. Heterogeneity within the MSC population further complicates matters, necessitating a comprehensive understanding and management of these diverse cell subsets to achieve consistent and predictable therapeutic outcomes. Additionally, the differentiation potential of MSCs can be influenced by various factors, requiring strategies to optimize their ability to differentiate into specific cell types for effective tissue repair. Furthermore, MSCs’ migratory capacity, a key factor in their regenerative effects, can be impacted by inflammation, age, and disease conditions, prompting the need for improved delivery methods or enhanced migratory capabilities to better target and repair injured tissues. Recent advancements in regenerative medicine have shed light on an intriguing alternative approach to exploit the therapeutic attributes of MSCs without utilizing the cells themselves to avoid potential immunocompatibility and other issues of cell-based therapies. This novel strategy involves utilizing MSC-derived exosomes for cell-free therapy^7,8^. Exosomes are nanosized extracellular vesicles secreted by cells and harbor diverse bioactive molecules, including cytokines and growth factors, signaling lipids, mRNAs, and miRNAs^9^. These MSC-derived exosomes can act as potent intercellular communicators, modulating cellular behaviors and facilitating tissue repair^10^. MSCs can be isolated from various tissues in the body, including bone marrow (BM), adipose tissue, connective tissue, umbilical cord, and dental pulp, among others. The tissue of origin can significantly influence the characteristics and therapeutic potential of MSC-derived exosomes^11^. However, it remains largely unknown whether these distinctions primarily arise from variations in the microenvironment between different tissue sources or if the tissue of origin itself plays a critical role in priming the therapeutic potential of exosomes. Particularly, the question of whether manipulating the MSC microenvironment can influence exosome content and therapeutic potential remains largely unexplored.

In our previous studies, we have demonstrated that macrophages, when treated with MSC-derived exosomes, enhanced the healing of Achilles tendons and ligaments in rodent models^4^. However, the macrophage-based treatment still confronts the same challenges encountered with other cell therapies (e.g., immunocompatibility, effectiveness for the cell delivery and cell storage). Our recent study has shown that exosomes derived from MSCs, when primed with the TLR-4 agonist LPS, exhibited improved efficacy in treating hematopoietic acute radiation syndrome^12^. These findings affirm that MSC exosomes play a role in tissue repair and that external priming of MSCs with pro-inflammatory mediators can alter their potency. Yet, it remains uncertain how priming MSCs influences the contents of exosomes, subsequently impacting the healing process.

In this study, we utilized a human BM MSC-derived exosomes *in vitro* system and a rat ligament injury *in vivo* model to address three fundamental questions: (a) whether the exosomal RNA content is determined by their parent cells; (b) whether the exosomal RNA content is customizable via perturbation of their parent cells’ microenvironment; (c) and how the alternation of RNA content in MSCs-derived exosomes affect the healing outcome in a ligament injury animal model. We found the context-dependent correlations between exosomal microRNAs (miRNA), and messenger RNAs (mRNA) to the RNAs in their parent cells. Specifically, under a normal (native) condition, there is a moderate correlation between exosomal RNAs and parental (MSCs) RNAs. The correlation increased significantly, especially for microRNAs, once the parent cells underwent microenvironment changes (e.g., per-condition MSCs using the proinflammatory cytokine TNF-α, or a synthetic Toll-like receptor 4 agonist (CRX-527)^13^, respectively). We showed that MSCs-derived exosomes can significantly increase ligaments’ max load and stiffness during the healing process based on a rat ligament injury *in vivo* model. We also demonstrated that the RNA content in exosomes is not only customizable via the perturbation microenvironment of the parent cells but also affects the ligament healing efficiency and mechanical properties. We further unraveled the molecular mechanisms underlying the altered healing capacity after treatment with native and pre-conditioned MSC-exosomes. Notably, we observed a direct connection between the changes in exosomal microRNA abundance and subsequent alterations in the expression of microRNA targets upon *in vivo* exosome delivery in the ligament. These insights shed light on the intricate communication between cells and their exosomes, providing a deeper understanding of the regulatory roles of exosomal RNAs in tissue repair and regeneration.

## Results

### Exosomal mRNAs from MSCs show enrichment in diverse biological functions but with limited overlapping with exosomal proteins

Although we have known that exosome cargoes are known to be enriched in proteins and small RNAs^14^, the extent to which exosomes are enriched in polyadenylated mRNAs and their potential functional relevance remains largely unknown. We performed RNA-seq on MSCs derived exosomes and identified 4,238 genes that exhibited significant expression in MSC-exosomes, with average normalized TPM values among replicates exceeding 1 (see **Supplementary Table S1**). These polyadenylated mRNAs were found to be enriched in diverse biological functions (see **Supplementary Table S2**). Interestingly, the top eight enriched pathways primarily pertained to the translation machinery (**Figure 1**). We conducted proteomics profiling of MSC-derived exosomal proteins to explore further the relationship between the abundance of exosomal RNAs and exosomal proteins. Surprisingly, our data indicated no correlation between the abundance of exosomal proteins and that of exosomal mRNA (**Supplementary Figure S1A**; Spearman’s rank correlation coefficient Rho = 0.018). We extended this analysis by treating MSCs with CRX-527 (referred to as CRX), a synthetic TLR4 agonist known for its potential as a vaccine adjuvant or for inducing anti-tumor immunity similar to G100. Consistent with native (untreated conditions), we did not observe any correlation between exosomal protein abundance and exosomal mRNA abundance in pre-conditioned MSCs derived exosomes (**Supplementary Figure S1B**; Rho = 0.055). Thus, our results indicate that exosomal mRNA abundance may be independent of exosomal protein abundance, regardless of the specific microenvironmental context.

**Figure 1:**
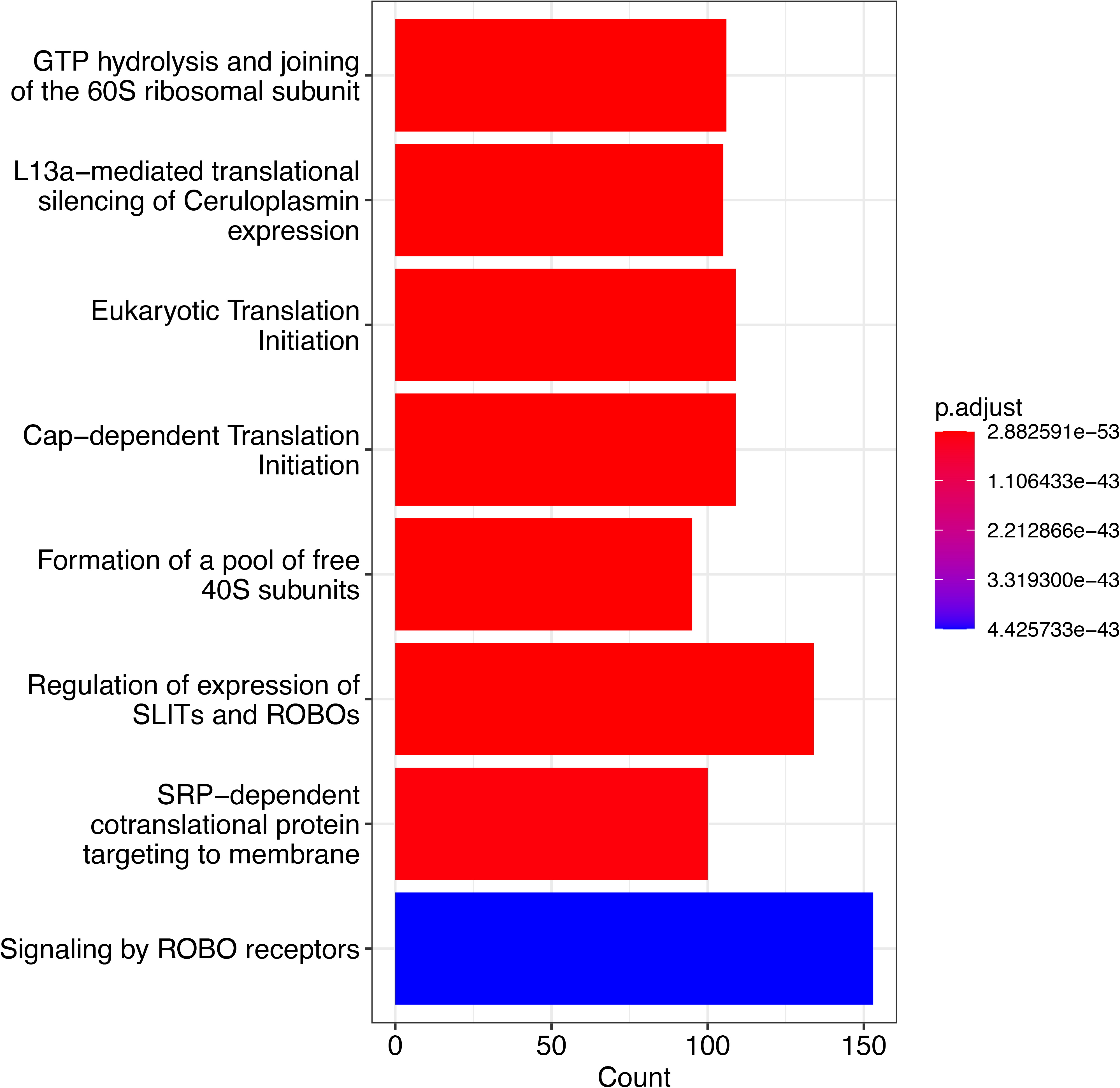
Top eight enriched pathways based on Benjamini-Hochberg (BH) adjusted p-values for MSC-exosome encapsulated mRNAs (average TPM > 1 for replicates).

### Microenvironmental perturbations of parent cells can influence the content of exosomal mRNAs

To investigate the impact of microenvironmental perturbation on exosomal mRNAs, we treated MSCs with CRX-527 and compared the levels of exosomal mRNAs between the treated and untreated cells. We identified 1,915 protein-coding genes that exhibited differential expression upon CRX-527 treatment (FDR<5% and >2 fold-change of normalized read counts; see the Methods section for details). This finding suggests that exosomal mRNAs can be modulated by their parent cells’ microenvironmental perturbations. We observed a significant enrichment of the “inflammatory response” gene ontology term (GO:0002544) among the CRX-527 induced up-regulated exosomal mRNAs (**Figure 2**; adjusted P-value = 3.8E-4). Interestingly, previous animal studies have shown that CRX-527 can elevate levels of cytokines and chemokines, thereby enhancing the host’s innate immune response and combating bacterial infections like Burkholderia pseudomallei^15^. This implies that the exosomes can potentially transport transcriptomic response signals from the parent cells under stimulating conditions. In addition to the inflammatory response GO term, the up-regulated exosomal mRNAs enriched various biological functions. These functions include, but are not limited to, positive regulation of fibroblast migration (GO:0010763; adjusted P-value = 6.2E-12), regulation of intrinsic apoptotic signaling pathway (GO:2001242; adjusted P-value = 1.4E-7), and plasminogen activation (GO:0031639; adjusted P-value = 8.9E-7). These results suggest that exosomal mRNAs, originating from the microenvironmental perturbation of their parent cells, may play a complex role in modulating immune response.

**Figure 2:**
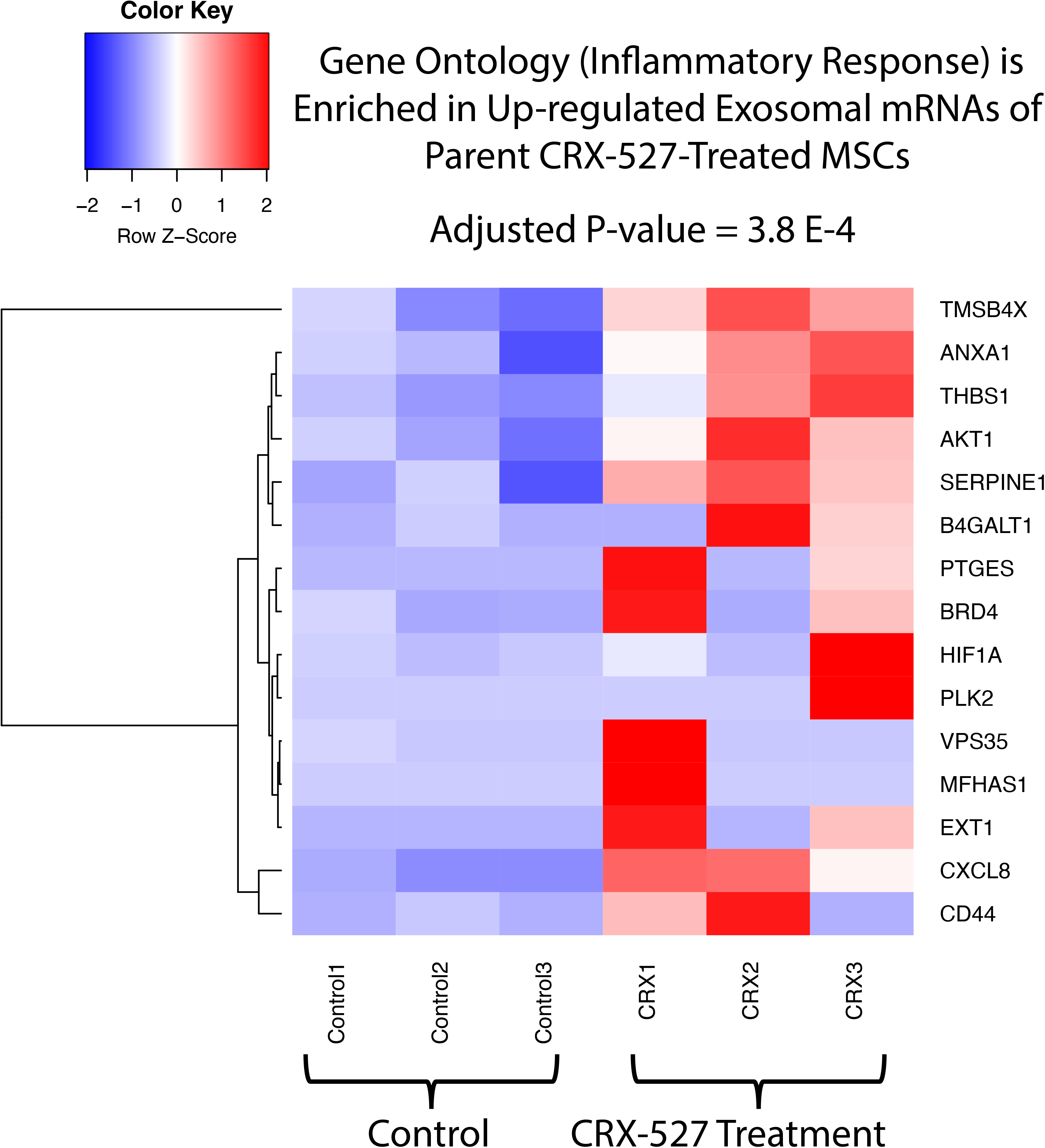
The illustration of the enrichment of up-regulated mRNAs in the inflammatory response gene ontology (GO) term. The heatmap displays the corresponding mRNA expression levels.

### Distinct microenvironmental perturbations of parent cells lead to the generation of unique exosomal mRNA contents

To determine whether changes in exosomal mRNA content are specific to particular microenvironmental perturbations or represent a general response, we compared the effects of two different treatments on parent cells (MSCs): tumor necrosis factor (TNF-α) and CRX-527. TNF-α is an adipokine known to communicate with various organs, including the brain, liver, muscle, immune system, and adipose tissue itself. It has been linked to insulin resistance, obesity-induced type 2 diabetes, fever induction, apoptotic cell death, cachexia, and inflammation. We sought to assess the extent of overlap in exosomal mRNA content changes between these two microenvironmental perturbations (TNF-α versus CRX-527). We observe a moderate overlap in the differentially expressed exosomal mRNAs between the two treatments (**Figure 3**). For instance, out of 180 exosomal mRNAs up-regulated in response to CRX-527 treatment, only 11 (6.1%) were up-regulated in response to TNF-α treatment. Regarding down-regulated genes, 857 out of 1,735 (49.4%) exosomal mRNAs affected by CRX-527 treatment overlapped with those affected by TNF-α treatment. Furthermore, the mRNA content changes associated with each MSC parent cell perturbation were found to be enriched in distinct biological functions (**Figure 3**). This observation suggests that exosomal mRNA content and their potential functions depend on the specific microenvironment of the parent cells.

**Figure 3:**
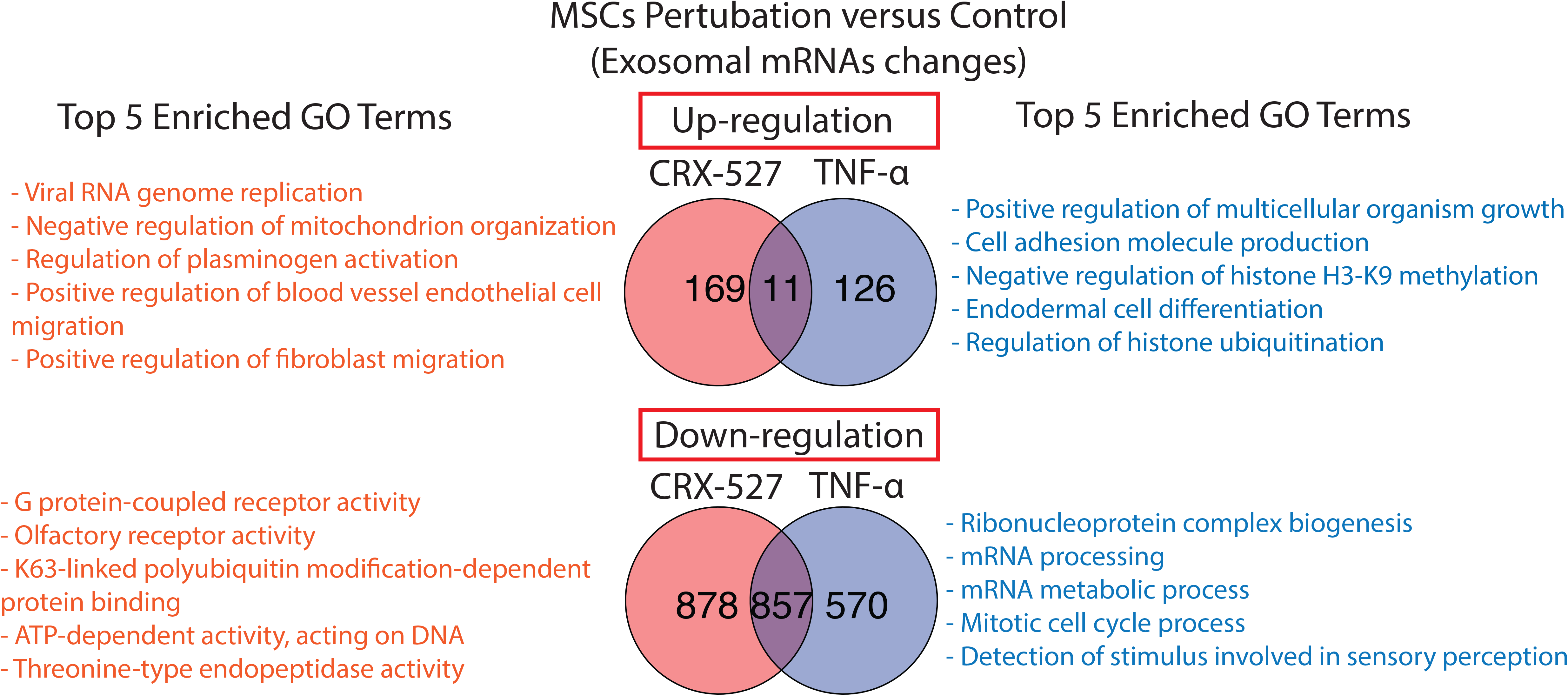
Comparison of mRNA content changes in different microenvironmental perturbations of the parent cells. The Venn diagram shows the overlapping up-or down-regulated mRNAs between CRX-527 and TNF-α treatments. The top 5 enriched Gene Ontology (GO) terms based on adjusted P-values for each perturbation are also shown.

### Microenvironmental perturbations of the parent cells can lead to the modulation of exosomal microRNA content

Recent evidence suggests that loading microRNAs into exosomes is a selective and regulated process rather than a random occurrence^16^. We aim to determine whether the parent cells’ microenvironmental perturbations can influence microRNA content loading into exosomes. To achieve this, we performed small RNA sequencing to profile short RNA fragments in exosomes derived from native MSCs, CRX-527, and TNF-α pre-conditioned MSCs. We identified 31 differentially expressed exosomal microRNAs in response to CRX-527 treatment (**Figure 4A**) and 23 differentially expressed exosomal microRNAs in response to TNF-α treatment in the parent cells (**Figure 4B**), respectively. Notably, none of these differentially expressed exosomal microRNAs overlapped between the CRX-527 and TNF-α treatments, suggesting that the content of exosomal microRNA is influenced by the specific microenvironment of the parent cells. We observed that “macrophage differentiation” (**Figure 4D**) is the top enriched gene ontology (GO) terms based on enrichment p-value for the microRNA targets of TNF-α pre-conditioned MSCs derived exosomes. TNF-α is a pro-inflammatory cytokine known to induce M1-like polarization of macrophages. Enriching the “macrophage differentiation” GO term suggests that as molecular cargo, exosomes may transport specific microRNAs to initiate monocyte differentiation into macrophages under pro-inflammatory stimuli such as TNF-α perturbation. **Figures 4C and 4D** present the top three enriched GO terms based on the best P-values for the CRX-527 and TNF-α perturbations, respectively. These findings highlight the broad functional impact of the microRNA changes observed.

**Figure 4:**
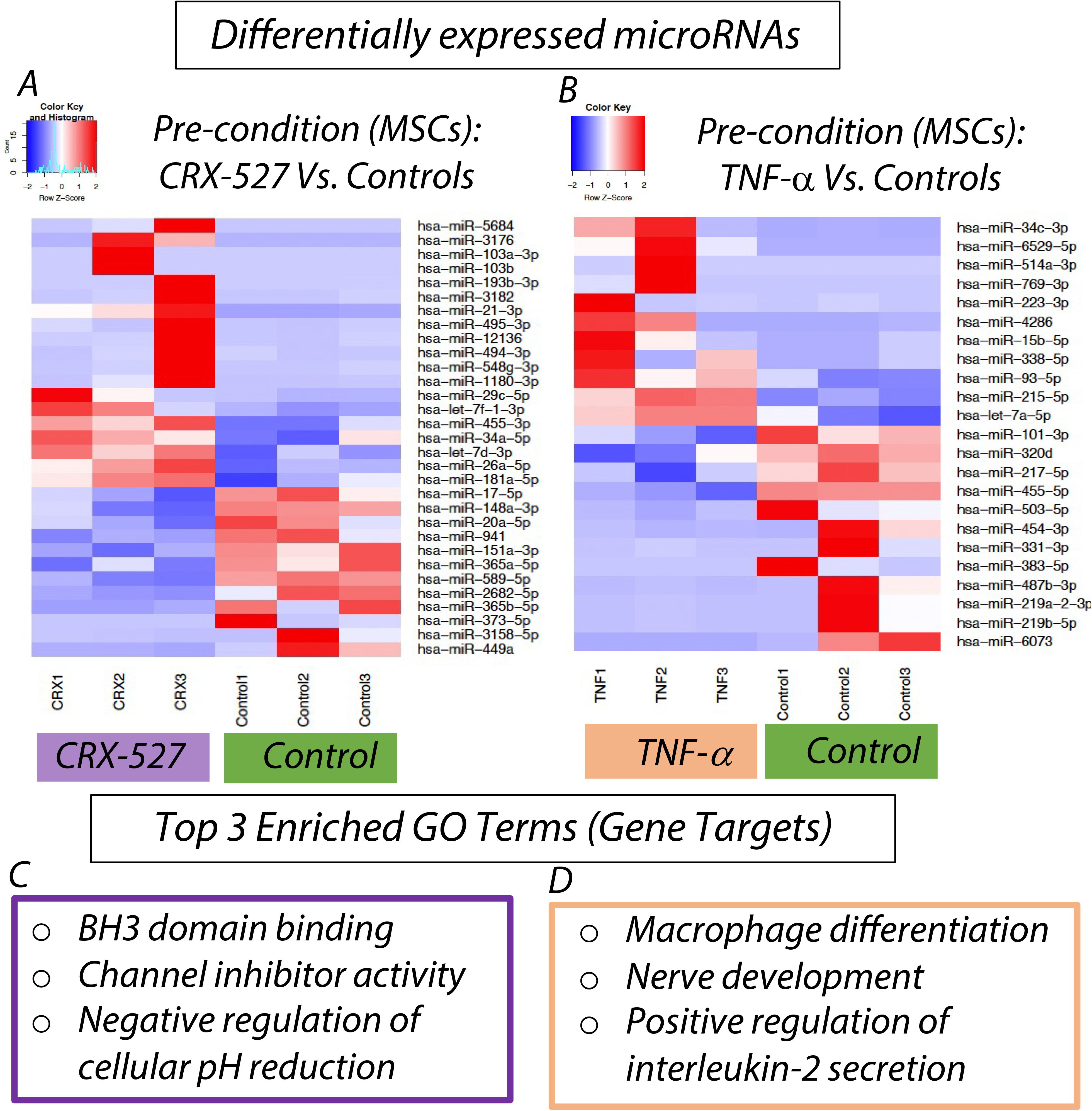
Exosomal microRNA levels are influenced by microenvironmental perturbations of the parent cells. (A) Differentially expressed exosomal microRNAs resulting from CRX-527 treatment in the parent cells (B) Differentially expressed exosomal microRNAs resulting from TNF-α treatment in the parent cells. The top three enriched gene ontology (GO) terms based on the best enrichment p-value for the microRNA targets are shown in Figure 4C and Figure 4D, respectively.

### Correlations in RNA abundance between parent cells and exosomes

We further conducted mRNA and small RNA sequencing on parent cells (MSCs) to investigate whether RNA abundance is correlated between parent cells and exosomes. We observed moderate correlations between exosomes and their parent cells under unperturbed conditions, with Spearman’s Rank correlation coefficients of 0.46 for microRNAs and 0.57 for mRNAs, respectively (**Figure 5**). This suggests that a subset of exosomal RNAs indeed represents the RNA abundance in their parent cells. These correlations increased when parent cells were exposed to microenvironmental stimuli, such as CRX-527 and TNF-α treatment. Notably, this effect was more pronounced for microRNAs (**Figure 5**). We observe that the Spearman’s Rank correlation coefficient (Rho) between exosomal and parent cell microRNA abundance increased from 0.46 (under native conditions) to 0.78 and 0.76 for CRX-527 and TNF-α pre-conditioned MSCs, respectively. This indicates that microenvironmental stimuli synchronize the microRNA profiles of exosomes with those of their parent cells. A similar trend was observed for mRNAs, albeit less prominently (**Figure 5**). These findings demonstrate that exosomal RNAs are not simply a random subset of RNAs derived from their parent cells. Rather, the selection of exosomal RNAs is regulated and depends on the transcriptomic status of their parent cells, including microenvironmental stimuli.

**Figure 5:**
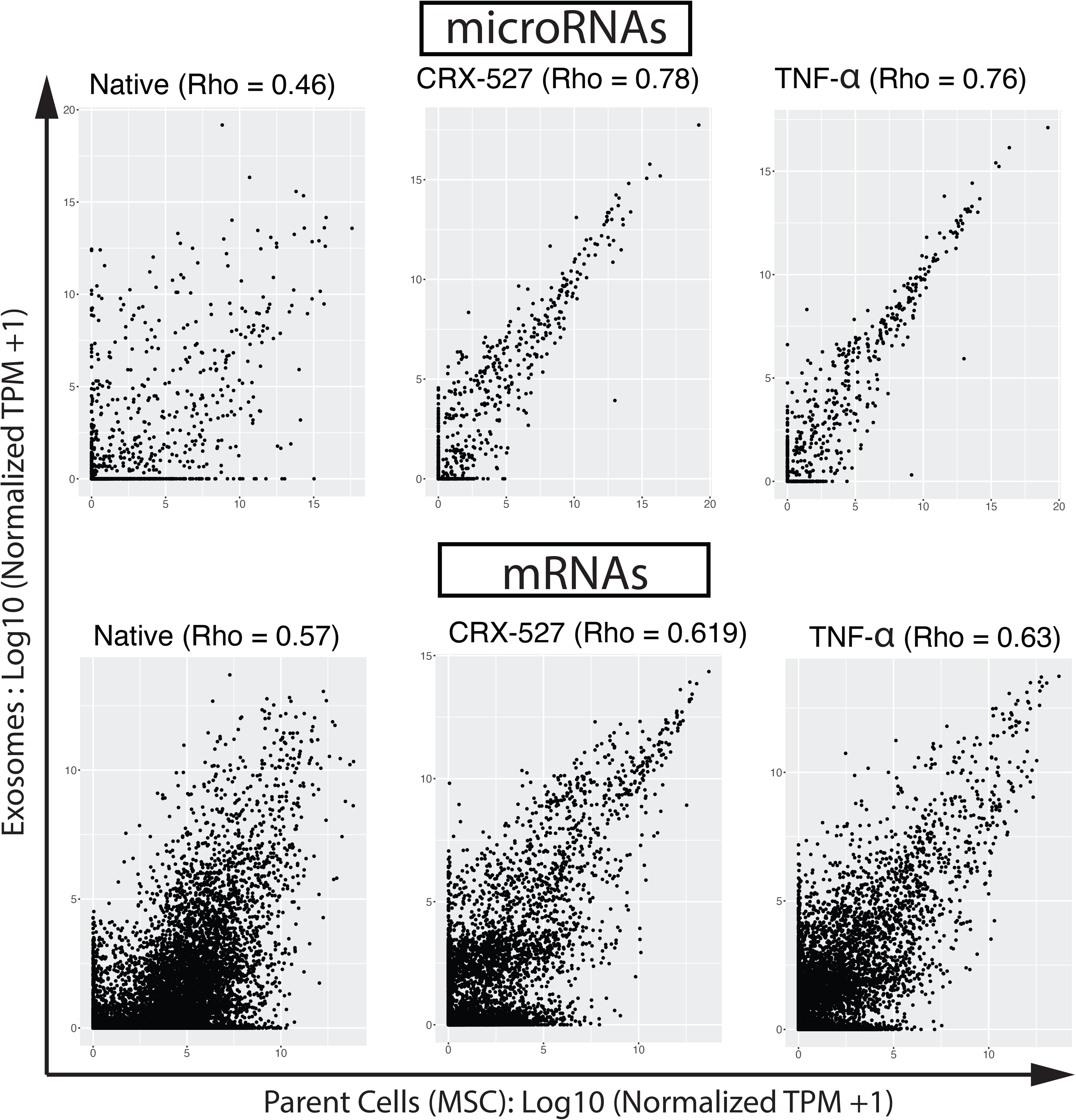
Context-dependent correlations between exosomal and parental RNA abundance. The Spearman’s rank correlation coefficient (Rho) is indicated.

### Immune pre-conditioned MSCs-exosomes affect the mechanical properties of the ligament healing

We asked whether treatment with exosomes or exosomes from either CRX-527 or TNF-α pre-conditioned MSCs can affect the mechanical properties of ligament healing. We surgically transected ligaments in rats to induce injury and then applied exosome treatment within the first three days post-injury. The goal was to compare the mechanical properties on day 14 between unstimulated MSC-exosomes and those preconditioned with TNF-α and CRX-527 (See Methods). We observed that treatment with exosomes from unstimulated (native) MSCs significantly increased max load (P=0.038), and stiffness (P=0.046) compared to PBS (untreated control) based on Wilcoxon Rank-Sum test (two-sided) (**Figure 6**), suggesting MSC-exosomes treatment improves mechanical properties of ligament healing. We further compared whether pro-inflammatory factors (CRX-527 and TNF-α) perturbed microenvironments in parent cells can affect the mechanical properties of exosome treatment. As shown in **Figure 6**, both CRX-527 and TNF-α preconditioned MSC-exosomes induced changes in the mechanical properties of the ligament, including max load, max stress, stiffness, and modulus, compared to native MSC-derived exosomes. This suggests that the microenvironment of the parent cells can modify the mechanical properties of the ligament following exosome treatment.

**Figure 6.**
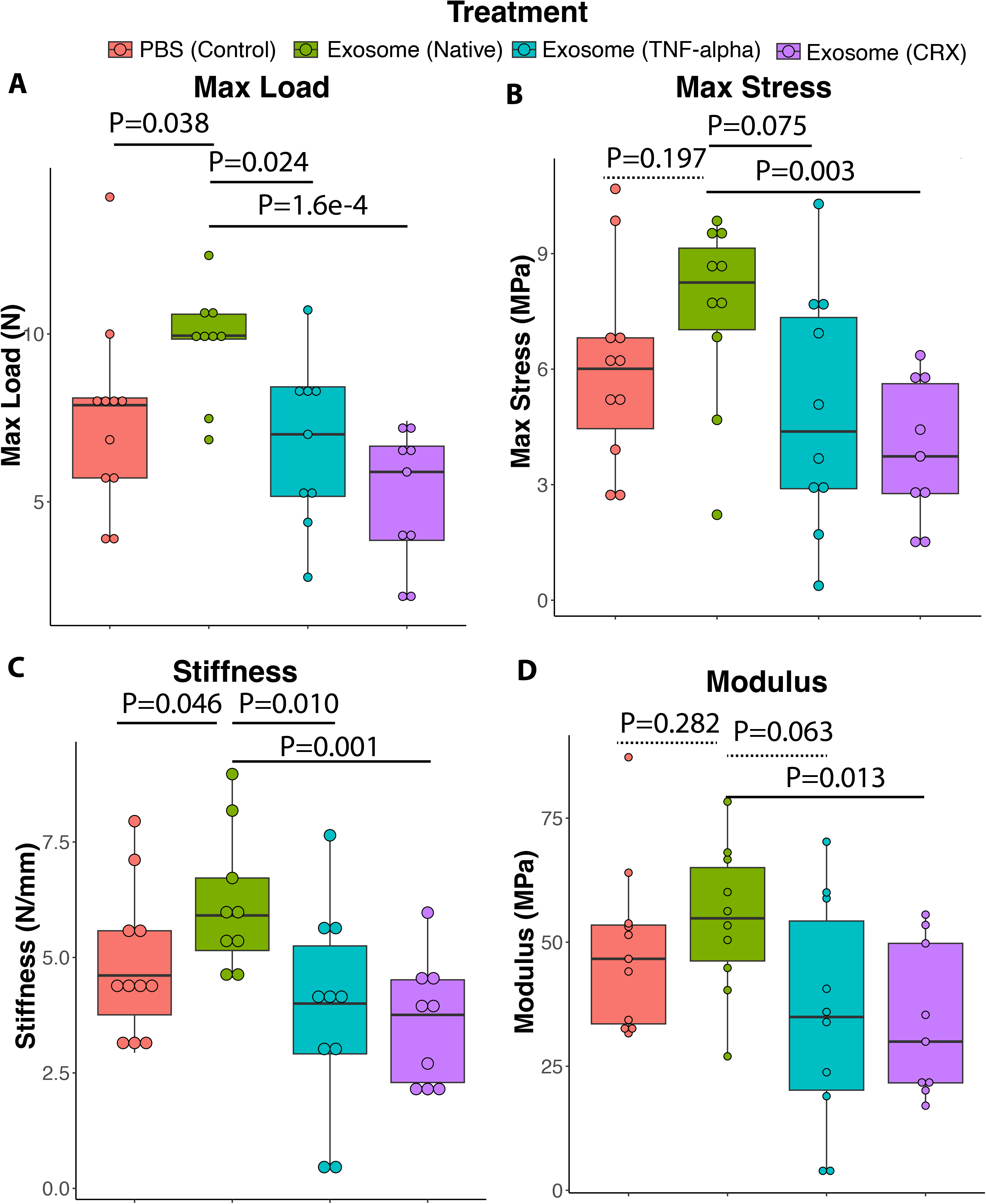
Mechanical properties of the healing ligament after treatment with native and pre-conditioned MSC-derived exosomes. Injured rat ligaments were treated with exosomes at early post-injury days (≤ 3 days). The mechanical properties from four types of exosomes (PBS-control, MSC-derived native exosome, TNF-α and CRX-527 preconditioned MSCs-derived exosomes) treatment were compared. (A) Max load; (B) Max stress; (C) Stiffness; (D) Modulus. The P-values (two-sided) of Wilcoxon Rank-Sum test for native exosome versus PBS-control and native exosome versus pre-conditioned exosomes are indicated. The solid lines indicate P<0.05, while the dashed lines indicate Pý0.05.

### Molecular mechanisms underlying the altered healing capacity upon treatment with native and pre-conditioned MSC-exosomes

To investigate the molecular mechanisms underlying the altered ligament mechanical properties influenced by MSC-exosomes, we conducted RNA-seq analysis (mRNA sequencing on post-injury day 14) on rat ligaments treated with native MSC-exosomes, pre-conditioned MSC-exosomes, and PBS controls (considered as untreated samples representing default healing). We identified 163 differentially expressed genes (DEGs) (fold change > 2 and FDR < 5%) in ligaments between native MSC-exosome treatment and PBS controls. The up-regulated genes were significantly enriched in multiple Gene Ontology (GO) terms associated with tissue regeneration, such as muscle contraction (adjusted P-value = 7.65E-6), positive regulation of epithelial morphogenesis (adjusted P-value = 0.001), and regulation of axonogenesis (adjusted P-value = 0.02). The top 15 most enriched GO terms for up-regulated genes based on adjusted P-values are presented in **Figure 7A**, while **Figure 7B** provides an example of the specific up-regulated genes enriched in muscle contraction terms.

**Figure 7:**
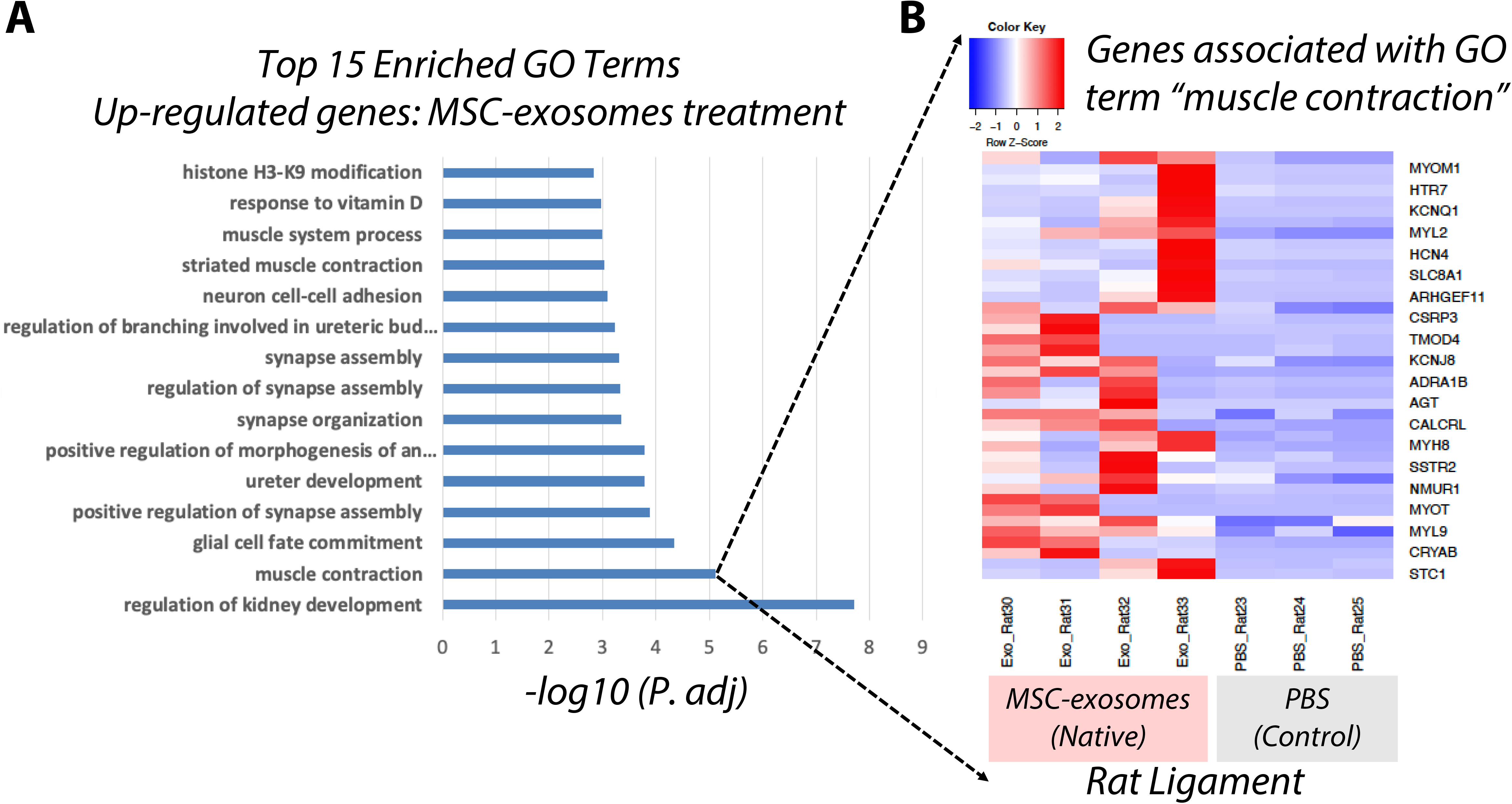
Enriched GO terms for MSC-exosome treatment induced up-regulated genes in rat ligament healing model. (A) Top 15 most enriched GO terms based on Benjamini-Hochberg (BH) adjusted P-values. (B) Up-regulated genes related to muscle contraction GO term in rat ligament induced by MSC-exosomes treatment.

Additionally, a wide range of GO terms were enriched in down-regulated genes in ligaments due to MSC-exosome treatment, including bone development (adjusted P-value = 0.006) and regulation of angiogenesis (adjusted P-value = 0.007), highlighting the complexity of the impact of MSC-exosome treatment on ligament healing. Detailed information on the enriched GO terms for up-regulated and down-regulated genes in MSC-exosome versus PBS control comparisons can be found in **Supplementary Table S3**.

Furthermore, we compared the transcriptomic responses of ligaments between pre-conditioned MSC-derived exosomes and native MSC-derived exosomes. We identified 163 and 286 differentially expressed genes for pre-conditioned CRX-527 and TNF-α exosomes compared to native exosomes, respectively. Among the top enriched terms for up-regulated genes in TNF-α pre-conditioned MSC-exosomes compared to native MSC-exosomes were GO terms related to the regulation of immune cells, such as regulation of T cell chemotaxis (**Supplementary Table S4**). In contrast, the up-regulated genes in ligaments pre-conditioned with CRX-527, compared to native MSC-exosomes treatment, were enriched in terms such as peptide hormone processing and nucleoside biosynthetic process (**Supplementary Table S5**), which are not related to immune cells. These findings indicate that the microenvironment of the parent cells can modulate the tissue *in vivo* response to exosome treatment.

### A connection between the alteration in exosomal microRNA abundance and the subsequent changes in the expression of microRNA targets induced by *in vivo* exosome delivery in ligaments

Since the abundance of exosomal microRNAs can be influenced by changes in the microenvironment after pre-conditioning of their parent cells (**Figure 4**), we conducted a microRNA-target enrichment analysis to investigate whether these changes in exosomal microRNA abundance directly impact the expression of downstream mRNA targets in rat medial collateral ligaments (MCLs). Because microRNAs typically regulate the expression of their targets by down-regulating them, we examined whether the targets of up-regulated or down-regulated exosomal microRNAs were more likely to be down-regulated or up-regulated, respectively, after *in vivo* exosome delivery. For each differentially expressed exosomal microRNA (e.g., exosomes from CRX-527 or TNF-α pre-conditioned MSCs versus native MSCs), we compared the ratio of microRNA targets among genes that were differentially expressed (> 2-fold changes & FDR < 5%) versus genes in the background (control group; < 1-fold change & FDR = 1) (**See Methods for details**). Among the differentially expressed exosomal microRNAs (pre-conditioned by CRX-527 and TNF-α versus native MSCs; Figure 4), nine of them (**Figure 8**) exhibited a statistically significant enrichment of microRNA targets (P-value < 0.05, Fisher’s Exact Test) among genes that were differentially expressed in response to the corresponding exosome treatment (pre-conditioned by CRX-527 and TNF-α versus native MSCs).

**Figure 8:**
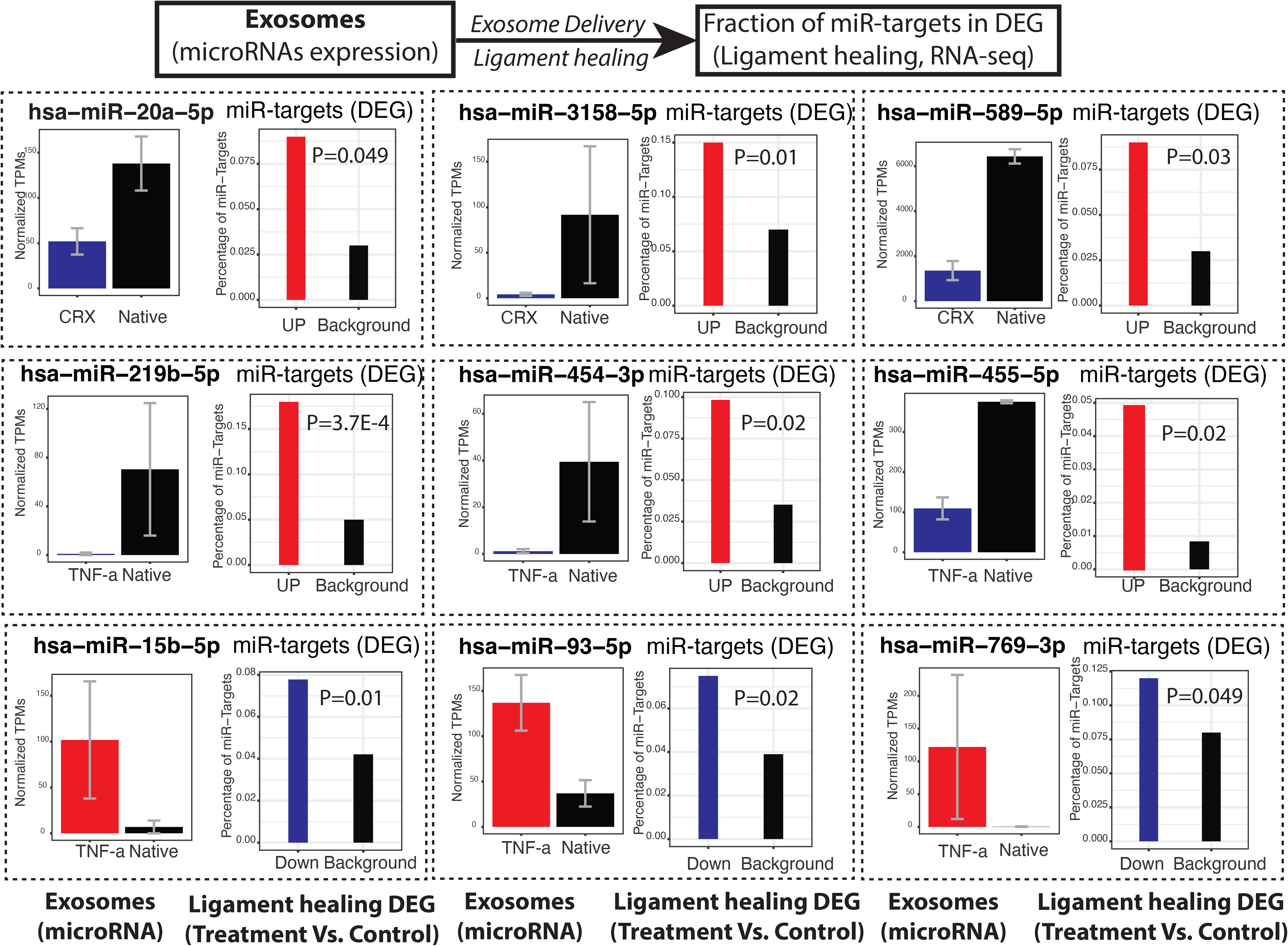
A connection between differentially expressed exosomal microRNAs and corresponding enriched microRNA-targets in response to exosome treatment. For each figure panel (dashed box), the left figure is the exosomal microRNA which shows differential pattern (Up-regulation or Down-regulation; > 2-fold changes & FDR <5%) between pre-conditioned (e.g., CRX-527 or TNF-α treatment) and native (untreated) MSCs. The right figure in each dashed box is the percentage of microRNA-targets in differentially expressed mRNAs (Down-regulated or Up-regulated) (> 2-fold changes & FDR <5%) and the background mRNAs (control; < 1-fold change & FDR=1). The statistical significance of each microRNA-target enrichment analysis (right figure in each dashed box) is calculated by Fisher’s exact test (2×2 contingency table: the number of DEGs (mRNAs; Up or Down) which are overlapped with corresponding microRNA targets in response to exosome (with and without pre-conditioned MSCs) treatment versus the number of background mRNAs (equivalent expressions) which are also overlapped with corresponding microRNA targets.

For instance, as illustrated in **Figure 8**, microRNA (miR-20a-5p) is down-regulated in exosomes derived from CRX-527 pre-conditioned MSCs (referred to as CRX-527-exosomes) compared to native MSC exosomes. After delivering these exosomes to rat ligaments, the up-regulated mRNAs in ligaments treated with CRX-527-exosomes, compared to native MSC-exosomes, were found to be statistically significantly enriched in miR-20a-5p targets. This indicates that the microRNA changes induced by pre-conditioned parent cells can contribute to corresponding changes in microRNA target expression in response to exosome treatment *in vivo*. It also reveals a mechanism whereby modulating the microenvironment of parent cells can influence the microRNA content of exosome cargo and subsequently impact the transcriptomic response to *in vivo* exosome treatment.

## Discussion

The present study reveals the intricate interplay between MSCs, their exosomes, and tissue regeneration, with a particular focus on ligament injury repair. To our best knowledge, this is the first demonstration that alterations in the MSC microenvironment using inflammatory mediators lead to significant changes in the exosome cargo RNA content. Notably, these alternations in molecular cargo were observable when exosomes were administered *in vivo* to a rat ligament injury model, underscoring that the exosome cargo is potentially customizable.

Another major discovery in this study is the context-dependent correlation between RNA content in parent cells and exosomes. We noticed a significantly enhanced correlation when the microenvironment of the parent cells was changed, particularly for microRNAs. This finding indicates the potential of exosomes to convey environmental changes or stimuli information from their parent cells to other cells.

The outcomes of our study provide insights into the adaptability and the versatile role of MSC-derived exosomes in tissue regeneration. Specifically, our findings on the customizable nature of exosome cargo and the dynamic RNA content correlation between parent cells and exosomes lay the foundation for developing tailored therapeutic strategies to enhance healing efficiency and mechanical properties in tissue repair. Moreover, the established link between altered exosomal microRNA levels and changes in microRNA target expression in ligaments unveils novel avenues for manipulating cellular communication for regenerative purposes.

This link offers a more nuanced understanding of the interaction between MSCs, their exosomes, and the regeneration of ligament tissues. Although our study focused on early post-injury time points, revealing increased ligament max load and stiffness, we recognize the importance of treatment timing and duration on ligament healing. Future research will delve into optimizing the exosome-based treatment strategy, exploring different time points and durations to ascertain the most efficacious approach. The use of TNF-α and CRX-527 as pro-inflammatory mediators to pre-condition MSCs raises intriguing questions about the influence of alternative pre-conditioning agents. Future studies could explore the use of anti-inflammatory mediators or growth factors to understand their impact on exosome RNA content and the ensuing effects on tissue regeneration.

## Conclusion

This study elucidates the intricate relationship between MSCs, their derived exosomes, and their impact on tissue regeneration. The newfound knowledge on the customizable exosome cargo and the context-dependent RNA content correlation opens up new horizons for developing tailored MSC-derived exosome therapies. By building on these findings and addressing the outlined future research directions, we can further refine and enhance the application of MSC-derived exosomes in regenerative medicine, bringing us a step closer to unlocking their full therapeutic potential.

## Materials and methods

### Cell culture

All protocols were approved by the Health Sciences Institutional Review Board of University of Wisconsin-Madison, School of Medicine and Public Health. Bone marrow (BM) derived MSCs were obtained as remnants from filters after bone marrow harvest of normal healthy donors^17^. BM cells were recovered by rinsing the filters with PBS. Mononuclear cells were separated using Ficol-Paque Plus 1.073 (GE Healthcare Bio-Sciences, Piscataway, NJ). Red blood cells were lysed in ACK lysis buffer, and mononuclear cells were suspended in α– minimum essential medium supplemented with 10% fetal bovine serum (US origin, uncharacterized; Hyclone, Logan, UT), 1× nonessential amino acids, and 4 mM L-glutamine. Cells were cultured in 75-cm^2^ filter cap flasks. Attached cells (passage 0) were harvested via TrypLE cell dissociation enzyme (Invitrogen, Carlsbad CA) and then re-plated into new flasks as described previously^18,19^ Cells (passages 4-6) were confirmed to be MSCs using flow cytometry and the typical marker profile^20^. MSC cultures were used for exosome isolation.

### Isolation and characterization of exosomes from MSCs

MSCs were grown to confluence in 75-cm^2^ filter cap cell culture flasks, washed with PBS, and the media was replaced with StemPro MSC serum-free media (SFM) CTS (Gibco Life Technologies, Gaithersburg, MD). Cells were incubated 18-24 hours upon which the conditioned culture media (CM) was collected and exosomes were isolated by differential ultracentrifugation essentially as previously described.^21^ The CM was centrifuged using a Beckman Coulter Allegra^®^ X-15R centrifuge (2000 *g* at 4°C for 20 minutes) to remove any detached cells, apoptotic bodies, and cell debris. Clarified supernatant CM was then centrifuged in a Beckman Coulter Optima™ L-80XP Ultracentrifuge (100,000 *g* at 4°C for 2 hours) with a SW 28 rotor to pellet exosomes. The supernatant was carefully removed, and exosome pellets were re-suspended in PBS and pooled. Exosomes were characterized using a Thermo NanoDrop spectrophotometer for protein and RNA concentration. Particle diameter and concentration were evaluated via IZON qNano Nanoparticle Characterization instrument (Zen-bio Inc, Research Triangle Park, NC).

### Medial collateral healing model

This study was approved by the University of Wisconsin Institutional Animal Care and Use Committee (Protocol number: M005785). All surgeries were performed using isoflurane for anesthesia. All efforts were made to minimize suffering. A total of 20 adult male Wistar rats (300-370 g) were used as an animal model for ligament healing.^22,23^ A surgically transected rather than torn ligament was used as an experimental model to create a uniform defect to compare healing. Rats were anesthetized via isoflurane and subjected to bilateral MCL transection. A 1 cm incision was made over the medial aspect of each stifle. The MCL was exposed, and transected at its mid-point. The ligament was not surgically repaired. To contain the treatments, the muscle layer was first sutured, then treatments were administered directly over the injured MCLs using a 28g needle and 1 ml syringe. All the treatments were performed at early post-injury days ((3 days). Treatments (30 ul) were delivered bilaterally and included (a) PBS (serving as the injured control), (b) 5 x 10^6^ exosomes (Exosome), (c) 5 x 10^6^ exosomes from TNF-α-preconditioned MSCs (TNF), or (d) 5 x 10^6^ exosomes from CRX-preconditioned MSCs. At 3 days post-injury, the left MCL received an additional dose of treatments. On day 14, MCLs were collected and used for mechanical testing. Day 14 was chosen as a collection time as the ligament to measure the treatment effects during early tissue remodeling. We did not observe a dose-dependent effect in mechanical testing, and thus we combined the left (two-dose) and right MCL (one-dose) treatment data in the downstream analysis.

### Mechanical testing

In order to test the structural properties of the healing ligament after exosome treatment, day 14 MCLs were mechanically tested. MCLs were dissected, and the surrounding tissue was excised^24^. Each MCL was removed with the femoral and tibial insertion sites intact. Ligaments remained hydrated using PBS. MCL length, width, and thickness were measured using digital calipers at pre-load. Width and thickness measurements were obtained at the injury site. The cross-sectional area (assumed to be an ellipse) was then estimated. The femur-MCL-tibia complex was mounted in a custom testing bath and mechanical testing machine. A pre-load of 0.1 N was applied, a cross-section was measured, and each MCL was preconditioned (cyclically loaded to approximately 2% strain for 10 cycles). The ligament was pulled to failure at a rate of 1% strain per second. Failure force, failure stress, ligament stiffness, and Young’s modulus were measured/computed to determine post-treatment MCL mechanical behavior. Failure force was the highest load prior to the failure of the ligament, and Lagrangian stress was calculated by dividing the failure force by the initial cross-sectional area of the ligament. Stiffness was calculated by determining the slope of the most linear portion of the load-displacement curve. Young’s modulus was calculated by the slope of the linear portion of the stress-strain curve.

### mRNA sequencing (RNA-seq)

Total RNAs were isolated from Human MSCs and MSCs-derived exosomes, as well as rat medial collateral ligaments (MCLs) (post-injury day 14) using trizol (ThermoFisher #15596018) and chloroform phase separations followed by the RNeasy mini protocol (Qiagen #74106) with optional on-column Dnase digestion (Qiagen #79254). One hundred nanograms of total RNA were used to prepare sequencing libraries using the LM-Seq (Ligation Mediated Sequencing) protocol^25^. RNAs were selected using the NEB Next Poly A+ Isolation Kit (NEB #E7490S/L). Poly A+ fractions were eluted, primed, and fragmented for 7 minutes at 85°C. First stand cDNA synthesis was performed using SmartScribe Reverse Transcriptase (Takara Bio USA #639538), and RNA was removed. cDNA fragments were purified with Ampure XP beads (Beckman Coulter #A63881). The 5’ adapter was ligated, and 18 cycles of amplification were performed. These final indexed cDNA libraries were quantified, normalized, multiplexed, and run as single-end reads on the HiSeq 3000 (Illumina, San Diego, CA). RNA-seq reads were mapped to the reference genome (human genome for human MSCs and MSC-exosomes; rat genome for rat ligament *in vivo* model) and corresponding annotated protein-coding genes using Bowtie (v0.12.8) ^26^, allowing up to 2-mismatches. The gene expected read counts and Transcripts Per Million (TPM) were estimated by RSEM (v1.2.3) ^27^. The TPMs were further normalized by EBSeq ^28^ R package to correct the potential batch effect.

### Small RNA-seq (microRNAs)

Total RNAs were isolated using trizol (ThermoFisher #15596018) and chloroform phase separations followed by the RNeasy mini protocol (Qiagen #74106) with optional on-column DNase digestion (Qiagen #79254). One hundred nanograms of total RNA was used to prepare sequencing libraries using the NEBNext Multiplex Small RNA Library Prep Set for Illumina (NEB #E7560S). The 3’ adapter was ligated and the reverse transcription primer hybridized. The 5’ adapter was ligated and cDNA synthesis was performed using ProtoScript II Reverse Transcriptase. PCR amplification was performed with 15 cycles and purified with Qiaquick PCR Purification Kit (Qiagen # 28106). cDNA libraries were then size-selected using AMpure XP beads (Beckman Coulter #A63881). These final indexed cDNA libraries were quantified, normalized, multiplexed, and run as single-end reads on the HiSeq 3000 (Illumina, San Diego, CA). small RNA sequencing reads were timed to the first 15 bp, and then mapped to mapped to the human mature microRNAs (miRbase^29^) using Bowtie (v0.12.8) ^26^. The microRNA expected read counts and Transcripts Per Million (TPM) were estimated by RSEM ^27^.

### Differentially expressed mRNAs and microRNAs

We used the EBSeq package ^28^ to assess the probability of a gene (mRNAs) or microRNAs being differentially expressed between any two given conditions. We required that DEGs should have FDR<5% via EBSeq and >2 fold-change of median-by-ratio normalized read counts.

### MicroRNA targets prediction and enriched gene ontology (GO) terms

The miRNA targets were predicted via the TargetScan database^30^. The Gene ontology (GO) enrichment analysis for the microRNA targets was performed by the R package (‘allez’) ^31^. The *P*-values were further adjusted by Benjamini–Hochberg (BH) multiple-test correction ^32^. *P*-values <0.05 were considered statistically significant.

### Reactome pathway enrichment analysis

The Reactome pathway enrichment analysis for MSCs-exosome enriched mRNAs was performed by the “ReactomePA” R package^33^.

### MicroRNA target enrichment analysis

To investigate the relationship between exosomal microRNA abundance and their downstream mRNA targets in rat ligaments, we employed a 2×2 contingency table to calculate the statistical significance of this microRNA target enrichment analysis. The contingency table was designed based on the widely accepted notion that an up-regulation in microRNA usually corresponds to a down-regulation of its mRNA target and vice versa. we examined whether the targets of up-regulated or down-regulated exosomal microRNAs (CRX-527 or TNF-α pre-conditioned MSCs versus native MSCs) were more likely to be down-regulated or up-regulated, respectively, after in vivo exosome delivery.

Specifically, we calculated four numbers (namely a, b, c, and d) in the table (taking up-regulated microRNAs as an example):

(a) a: the number of instances where the down-regulated genes (mRNAs) in rat ligaments after in vivo exosome delivery are also the targets of up-regulated microRNA;
(b) b: the number of instances where the down-regulated genes (mRNAs) in rat ligaments after in vivo exosome delivery are not the targets of up-regulated microRNA;
(c) c: the number of instances where the background genes (|fold-change| < 1 and FDR = 1) are the targets of up-regulated microRNA;
(d) d: the number of instances where the background genes (|fold-change| < 1 and FDR = 1) are not the targets of up-regulated microRNA;

We calculated the p-values of Fisher’s exact test to interrogate whether the ratio of (a/b) is statistically greater than the ratio of (c/d), asking whether the down-regulated mRNAs in rat ligaments is significantly enriched in the targets of up-regulated exosomal microRNA.

A similar approach is also applied to calculate the statistical enrichment of whether up-regulated mRNAs in rat ligaments is significantly enriched in the targets of down-regulated exosomal microRNA.

### Nanoscale liquid chromatography coupled to tandem mass spectrometry (nano LC-MS/MS)

Nano LC-MS/MS was used to profile proteomics of parent MSCs and their MSCs-derived exosomes. Digests were desalted using 100µl Pierce® C18 Tips (Thermo Fisher Scientific) according to manufacturer protocol. Eluates in 70%:30%:0.1% Acetonitrile:Water:TFA acid (vol:vol) were dried to completion using speed vac and finally reconstituted for LC-MS/MS profiling. Peptides were analyzed on a newer generation Orbitrap Fusion™ Lumos™ Tribrid™ platform. Chromatography of peptides prior to mass spectral analysis was accomplished using capillary emitter column (PepMap® C18, 2µM, 100Å, 500 x 0.075mm, Thermo Fisher Scientific). NanoHPLC system delivered solvents A: 0.1% (v/v) formic acid, and B: 80% (v/v) acetonitrile, 0.1% (v/v) formic acid at 0.30 µL/min to load the peptides at 2% (v/v) B, followed by quick 2 minute gradient to 5% (v/v) B and gradual analytical gradient from 5% (v/v) B to 37.5% (v/v) B over 73 minutes when it concluded with rapid ramp to 95% (v/v) B for a 5minute flash-out. As peptides eluted from the HPLC-column/electrospray source survey MS scans were acquired in the Orbitrap with a resolution of 120,000 followed by HCD-type MS2 fragmentation into Ion Trap (32% collision energy) under ddMSnScan 1 second cycle time mode with peptides detected in the MS1 scan from 350 to 1600 m/z; redundancy was limited by dynamic exclusion and MIPS filter mode ON. Lumos acquired MS/MS data files were searched using Proteome Discoverer (ver. 2.5.0.400) Sequest HT search engine against Uniprot *Homo sapiens* database (UP000005640, 08/11/2020 download, 96,816 total sequences). Static cysteine carbamidomethylation, and variable methionine oxidation plus asparagine and glutamine deamidation, 2 tryptic miss-cleavages and peptide mass tolerances set at 10 ppm with fragment mass at 0.6 Da were selected. Peptide and protein identifications were accepted under strict 1% FDR cut-offs with high confidence XCorr thresholds of 1.9 for z=2 and 2.3 for z=3. Strict principles of parsimony were applied for protein grouping. Chromatograms were aligned for feature mapping, and ion intensities were used for precursor ion quantification using unique and razor peptides. Normalization was performed on the total peptide amount and scaling on all averages.

## Availability of data and materials

We have submitted the raw RNA-seq data and the gene expression data (TPMs and Mapping counts) to GEO (accession number: GSE245884).

## Acknowledgments

We thank Heather Hartwig-Stokes (MS, UW-Madison) for mechanical testing, Ellen M. Leiferman (DVM, UW-Madison) for student surgical training. Funding support: NIH/NHLBI (R0HL153721 to Christian M. Capitini and Peiman Hematti) and DARPA (AWD00001593 to Peng Jiang). We thank the support from the Center for Gene Regulation in Health and Disease (GRHD) at Cleveland State University and Prof. Anton Komar.

## Author contributions

- Concept and experiment design: CSC, PJ
- Exosome experiment: CSC, WM, RV
- Animal experiments: CSC, WM, RV
- Proteomics profiling: JK, CMC, and PH
- RNA-seq (mRNA and microRNAs) profiling: PJ
- Technical support: EM, YL, SY, CMC
- Data analysis: AP & PJ
- Manuscript writing: CSC, AP and PJ

## List of supplementary information

**Supplementary Table S1**: List of genes that exhibited significant expression in MSC-exosomes, with average normalized TPM values among replicates exceeding 1.

**Supplementary Table S2:** Enriched pathways for genes that exhibited significant expression in MSC-exosomes.

**Supplementary Table S3:** GO terms enriched in genes that are either up-regulated or down-regulated in rat ligaments treated with MSC-exosomes compared to the PBS control.

**Supplementary Table S4:** GO terms enriched in genes that are either up-regulated or down-regulated in rat ligaments treated with TNF-α pre-conditioned MSC-exosomes compared to the treatment using native MSC-exosomes.

**Supplementary Table S5:** GO terms enriched in genes that are either up-regulated or down-regulated in rat ligaments treated with CRX-527 pre-conditioned MSC-exosomes compared to the treatment using native MSC-exosomes.

**Supplementary Figure S1:** The correlations between the abundance of exosomal mRNAs and exosomal proteins. Spearman’s rank correlation coefficient (Rho) is indicated. (A) native exosome; (B) Exosome derived from CRX-527 pre-conditioned MSCs.

## Supplementary Figures

**Supplementary Figure S1:**
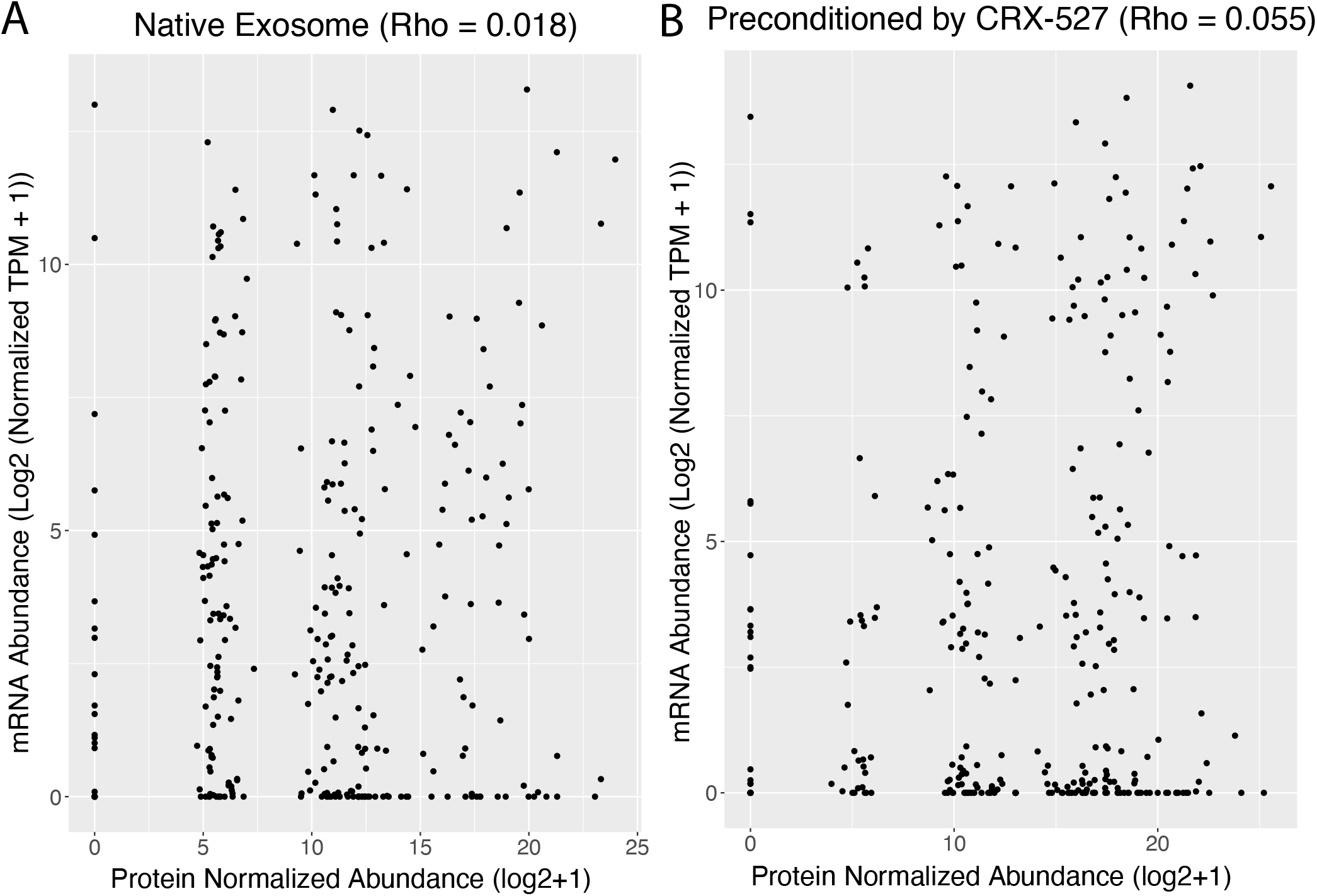
The correlations between the abundance of exosomal mRNAs and exosomal proteins. Spearman’s rank correlation coefficient (Rho) is indicated. (A) native exosome; (B) Exosome derived from CRH-527 pre-conditioned MSCs.

